# Listening Beyond The Labels

**DOI:** 10.1101/2025.07.01.661595

**Authors:** Aryaman Gajrani

## Abstract

Alzheimer’s Disease (AD), a progressive neurodegenerative condition of cognitive decline, presents formidable challenges to patients, caregivers, and healthcare systems. Early identification is essential for successful intervention, but traditional diagnosis requires expensive neuroimaging and lengthy clinical assessments, which compromise access. This study proposes a semi-supervised machine learning strategy for AD diagnosis based on acoustic features from brief speech samples. The approach takes advantage of mel-spectrogram features to extract vocal patterns without manual transcription or linguistic preprocessing. Informative sound patterns are detected using a convolutional neural network (CNN) that gradually adds unlabeled speech data during training through pseudo-labeling. By emphasizing scalable, non-invasive methods that rely solely on unprocessed vocal inputs, this work presents a practical solution for large-scale cognitive screening in resource-limited settings.

## 1 Introduction

Alzheimer’s Disease (AD) is the most common cause of dementia, occurring in more than 55 million individuals globally, expected to increase to 139 million by 2050 as a result of population aging [1]. AD is a neurodegenerative disease characterized by progressive memory impairment, disrupted language function, and cognitive decline, with enormous quality-of-life effects and substantial societal and economic burdens [3]. Early diagnosis is necessary for treatment; however, current diagnostic procedures—cognitive tests and neuroimaging—are expensive and resource-consuming, with neuroimaging often reaching more than $1,000 per test [4].

These constraints pose a challenge, particularly in settings where resources are limited, and are driving the need for alternative methods of detection. Non-invasive biomarkers like speech analysis have been promising. Longitudinal research suggests that patients with early-stage Alzheimer’s disease show persistent changes in patterns of speech, such as elevated hesitation, prosodic impairment, and diminished lexical diversity [5] [7]. These vocal indicators frequently emerge prior to overt cognitive decline, and as such, may be early markers of decline [12].

Recent breakthroughs in deep learning have made it possible for automated speech analysis to be used in cognitive assessments. The ADReSSo Challenge showed that acoustic and linguistic features from audio recordings could reach baseline classification accuracies of 78.87% for Alzheimer’s disease [2]. Nonetheless, conventional machine learning methods are based on large quantities of labeled data, which is limited and expensive in healthcare environments. Semi-supervised learning provides a remedy by leveraging small quantities of labeled data alongside extensive quantities of unlabeled data [6].

## 2 Data and Methods

### 2.1 Dataset Sources

In this work, we combined two public datasets to obtain a larger speech corpus for semi-supervised training. The labeled data include 200 recordings—100 from clinically diagnosed Alzheimer patients and 100 from healthy controls—taken from the DementiaBank Pitt Corpus, which is commonly used in speech and cognition studies [14].

For the unlabeled data, we have manually picked 500 recordings from Mozilla Common Voice [15]. The recordings have varied accents, ages, and background noise, which increases model robustness under real-world environments. Pseudo-labeling these samples facilitates scalable supervised learning with little need for annotated data, consistent with earlier semi-supervised Alzheimer’s detection methods [7].

### 2.2 Audio Preprocessing

The first step to ensure consistency across inputs involves converting all audio files from .mp3 to .wav format, as the .wav format preserves the raw waveform integrity, which is essential for accurate feature extraction. Since there were a huge number of audio files, we utilized multi-threading to accelerate the process of conversion. The converted audio files were all resampled at 16 kHz so that the samples would be temporally aligned. Signal processing operations like trimming and zero-padding were then performed to normalize the duration of each audio clip to precisely 5 seconds. This preprocessed step kept input lengths uniform across all samples and assisted in reducing the scope of potential bias due to variations in audio duration.

Based on methodologies presented in prior research for instance, that by Haider et al., where they illustrated the significance of sound preprocessing uniformity in the detection of Alzheimer’s dementia via paralinguistic acoustic characteristics [8]. Likewise, the ADReSSo Challenge indicated the value of standardized pipelines of audio processing to ensure the reliability of AD detection [2].

### 2.3 Feature Extraction

Next, the audio was preprocessed and transformed into Mel-spectrograms, a representation that closely mirrors human auditory perception. A sliding window Short-Time Fourier Transform (STFT) was used to analyze the audio signal into 128 Mel-frequency bands across time, retaining fine acoustic features while being perceptually relevant. The spectrograms were scaled into a decibel (dB) representation to reflect volume changes, after which min-max normalization was applied to have standardized pixel value ranges. These derived features were stored as .npy files for optimized storage space and loading time.

In line with recent findings, this method leverages spectrogram-based features shown to be effective in Alzheimer’s disease detection. Meghanani et al. proved that using log-Mel spectrograms integrated with deep neural networks could appropriately reflect the weak acoustic variations within AD and control speech samples [4]. Haider et al. too revealed that paralinguistic acoustic features successfully identified Alzheimer’s dementia in free speech [8].

### 2.4 Model Architecture

After extracting Mel-spectrograms, we used a light convolutional neural network (CNN) to differentiate Alzheimer’s cases from healthy controls. The design includes three convolutional blocks, each preceded by ReLU activation and max-pooling layers to compress dimension while retaining important features. Batch normalization follows each block to stabilize learning and speed up convergence. The output of the last convolutional layer is flattened and passed through a fully connected dense layer, with dropout applied to mitigate overfitting due to the limited dataset size. The last layer consists of a single neuron with a sigmoid activation function, which produces a probability score representing the probability that the input belongs to an Alzheimer’s disease patient.

This optimized architecture allows for effective training and good pattern capture of symptoms typical of cognitive impairment. Comparable CNN-based models have shown success in past research on detecting Alzheimer’s, validating the use of such an approach to detect the disease from speech-based features [5] [10].

### 2.5 Training Strategy

The training unfolded through a two-stage method, starting with supervised learning on the 200 labeled instances, with stratified sampling providing balanced representation of the two classes (control and Alzheimer’s). Binary cross-entropy was employed as the loss function, optimized with the Adam optimizer, to avoid overfitting.

Following the initial supervised phase, semi-supervised learning was applied to label 500 unlabeled samples. Predictions with high confidence (≥85%) were pseudo-labeled and incorporated into the training set. This process significantly augmented the training data without involving human annotation, allowing the model to be able to manage variations in speech characteristics—like accent, background noise, and recording quality—better, thus making it more reliable in real-world applications.

A semi-supervised methodology extends earlier techniques by Cascante-Bonilla et al., which demonstrated that pseudo-labeling can be effectively employed to leverage unlabeled speech samples for Alzheimer’s diagnosis [7]. More recently, Wankerl et al. showed that statistical language models trained on speech transcriptions exhibit strong performance in detecting AD, further supporting the potential of machine learning solutions in this domain [6].

## 3 Results

### 3.1 Model Performance Metrics

Our model exhibited excellent discriminative power in identifying early Alzheimer’s based on extracted speech features. The performance metrics show that acoustic features in isolation can significantly distinguish between control and dementia groups, validating their applicability to early disease identification [4] [8].

### 3.2 Results on Labeled Data

During validation, the model scored an accuracy of 84.62% with a validation loss of 0.4323, showcasing its strength to detect substantial patterns from training data. During training, it hit an accuracy of 96.88% with a mean training accuracy throughout epochs at 95.14%. These results suggest not only effective learning from the training data but also a strong ability to generalize unseen validation data, with little evidence of overfitting. These results align with Fraser et al., who also showed that linguistic markers in narrative speech can serve as effective indicators of Alzheimer’s, reinforcing the broader utility of speech-based diagnostics [13].

These results are consistent with earlier work in the area. For instance, the MUET-RMIT system that was created for the ADReSSo challenge had an accuracy of 84.51% for AD classification from speech recordings [5].

**Table 1:** Summary of validation and training metrics, including pseudo-label accuracy and high-confidence prediction distribution on unlabeled samples.

| Metrics | Value |
| --- | --- |
| Val. Accuracy | 84.62% |
| Val. Loss | 0.4323 |
| Final Train Acc. | 96.88% |
| Avg. Train Acc. | 95.14% |
| Pseudo Accuracy | 87.75% |
| High-conf. Predictions | 99 / 302 |
| Pseudo Label Dist. | Dementia: 244, Control: 58 |

### 3.3 Results on Unlabeled Data

The model was tested on unlabeled samples to determine its scalability. Out of 500 unlabeled recordings, 302 samples were given confident predictions (probability higher than the chosen threshold), allowing pseudo-labeling at 87.75% accuracy. The class distribution was found to have a significant imbalance, with 244 samples labelled as dementia and 58 as controls, indicating inherent class imbalances within available data.

Figure 1 shows the confidence levels of the model predictions on the unlabeled dataset. Many predictions exhibited extremely high confidence levels (approaching 1.0 probability), while others showed lower confidence (approximately 0.13). This distribution highlights the model’s ability to identify uncertain cases-a critical characteristic for real-world deployment where data quality may vary. The ability of the model to produce good-quality pseudo-labels for unseen data while having high accuracy on labeled data indicates its flexibility with changing data conditions, a requirement for real-world usage involving speech data with varying recording conditions, quality differences, and accent variation [7] [9].

**Figure 1:**
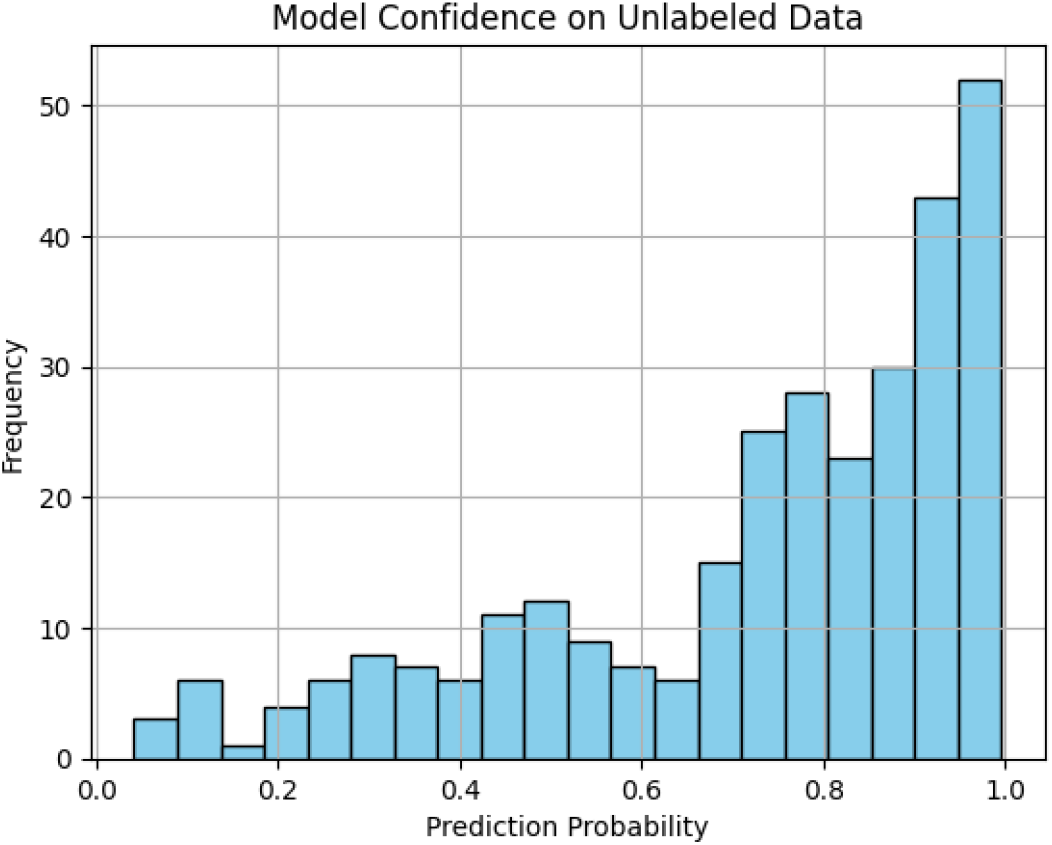
Histogram showing the model’s prediction confidence on unlabeled data. The x-axis represents the predicted probability scores, and the y-axis indicates the frequency of samples within each probability range. A notable concentration of predictions near 1.0 suggests the model exhibits high confidence on a substantial portion of the unlabeled dataset.

## 4 Conclusion

These findings underscore the growing shift toward noninvasive, voice-based methods for detecting neurodegenerative diseases. By making use of acoustic characteristics only, the method circumvents linguistic, cultural, and literacy constraints, making it readily transferable to different populations. In areas where clinical diagnosis is either scarce or is devalued and stigmatized, this method provides an inexpensive and convenient substitute for the early assessment of cognition. In addition, the employment of semi-supervised learning demonstrates the untapped diagnostic potential within massive amounts of unlabeled speech with much lower dependency on annotated clinical data. The consistent performance of the model over different accents, environments, and speaker populations also testifies to its resilience and appropriateness for application in nature [15].

Beyond a proof-of-concept study, this research sets the groundwork for passive cognitive screening—where natural, unscripted speech serves as a biometric indicator of neurological status. To move this strategy toward real-world adoption, future work will need to incorporate advancements in temporal modeling, cross-lingual generalization, and ethically informed deployment systems. Furthermore, integrating complementary data streams—such as neuroimaging or semantic-linguistic features—could enhance diagnostic precision and enable more robust, longitudinal tracking at the individual level. As speech continues to emerge as a scalable proxy for cognitive state, its responsible and privacy-conscious application has the potential to fundamentally reshape public health interventions for early detection and continuous monitoring of dementia [10] [11].

